# Emergence of stable motifs in consumer-resource communities

**DOI:** 10.1101/2020.09.03.280644

**Authors:** Priyanga Amarasekare, Ulrich Brose, Jonathan Chase, Tiffany Knight, Adam Clark

## Abstract

Understanding how and why complex communities can be stable has preoccupied ecologists for over a century. Data show that real communities tend to exhibit characteristic motifs and topologies. Despite a large body of theory investigating both ecological (niche partitioning) and evolutionary (speciation and extinction) mechanisms, a general explanation for why particular motifs are more common than others remains elusive. Here we develop a mechanistic framework that investigates the set of possible motifs that can emerge under minimal conditions of a nutrient-limited system with no external inputs, and no spatial heterogeneity. Focusing on consumer-resource communities structured by competition and predation, we find that the emergent motifs under these minimal conditions are vertical trophic chains that maximize energy transfer and biomass production. Not only are such motifs stable to perturbations of species’ abundances, but they are also robust to species additions and removals. Our findings provide a mechanistic explanation for why tri-trophic chains are overrepresented in real food webs. They suggest that, because they maximize energy transfer, and can emerge and persist under minimal conditions, vertical trophic chains may constitute the fundamental architecture of consumer-resource communities.

## Introduction

Elucidating the mechanisms that stabilize complex ecological communities is a central issue in community ecology. Recent years have seen network theory, the study of interactions and connectivity between the elements of a given system (Alon, 2003), playing a key role in this endeavor. Two network properties have been particularly important in studying complex communities. The first is modularity, the organization of entities (e.g., genes, cells, individuals, species) into subsets that interact more strongly amongst themselves than with other such groups (Milo et al., 2002; Kashtan and Alon, 2005). The second is the existence of motifs, small sets of recurring elements that appear at a higher frequency than expected by chance (Milo et al., 2002; Alon, 2007).

The preponderance of modularity in transcription, neuronal and signal transduction networks is attributed to three advantages. First, organization into discrete, individual units confers robustness (maintenance of structural and functional integrity Wagner (2005)) by containing the impact of perturbations within localized areas of the network. Second, such modules can be easily reconfigured to adjust to changing environments, thus increasing robustness (Lipson et al., 2002; Alon, 2003). Third, the modular organization can increase the efficiency of network activity (Kashtan and Alon, 2005).

Biological networks also exhibit characteristic motifs. Transcription networks in unicellular organisms (e.g., yeast and *E. coli*; Milo et al. (2002); Alon (2007)) are characterized by three network motifs: negative autoregulation (NAR), feed-forward loops (FFLs), and single input modules (SIMs). Signal transduction networks involved in protein modification (e.g., phosphorylation) exhibit bi-parallel (diamond) motifs, while neural networks exhibit FFLs, bi-parallels, and bi-fans (two source nodes directly cross-regulating two target nodes; Milo et al. (2002)).

Some of these same motifs are also observed in ecological communities (Fig. 1). Negative autoregulation is akin to intra-specific competition due to resource limitation. Feed-forward loops can resemble tri-trophic food chains or a closed loop with omnivory, with energy being transferred from lower to higher trophic levels. Single input modules (SIM) are akin to exploitative and apparent competition motifs in ecological networks, while the bi-parallel (diamond) motif represents the combination of exploitative and apparent competition. Indeed, previous studies have found that consumer-resource webs (e.g., predator-prey, plant-herbivore, host-parasite) exhibit four motifs: tri-trophic chain, omnivory, exploitative and apparent competition. Most studies find the tri-trophic chain to be more frequent than expected by chance (Milo et al., 2002; Camacho et al., 2007). Evidence for omnivory is more equivocal, with some studies finding it to be overrepresented (e.g., Kondoh (2008) and others finding it not (e.g., Milo et al. (2002); Bascompte and Melian (2005); Johnson et al. (2014)). Many studies find exploitative and apparent competition to be less frequent than expected by chance (Milo et al., 2002; Camacho et al., 2007). Only one study that we are aware of found the bi-parallel (diamond) motif to be overrepresented (Milo et al., 2002).

**Figure 1:**
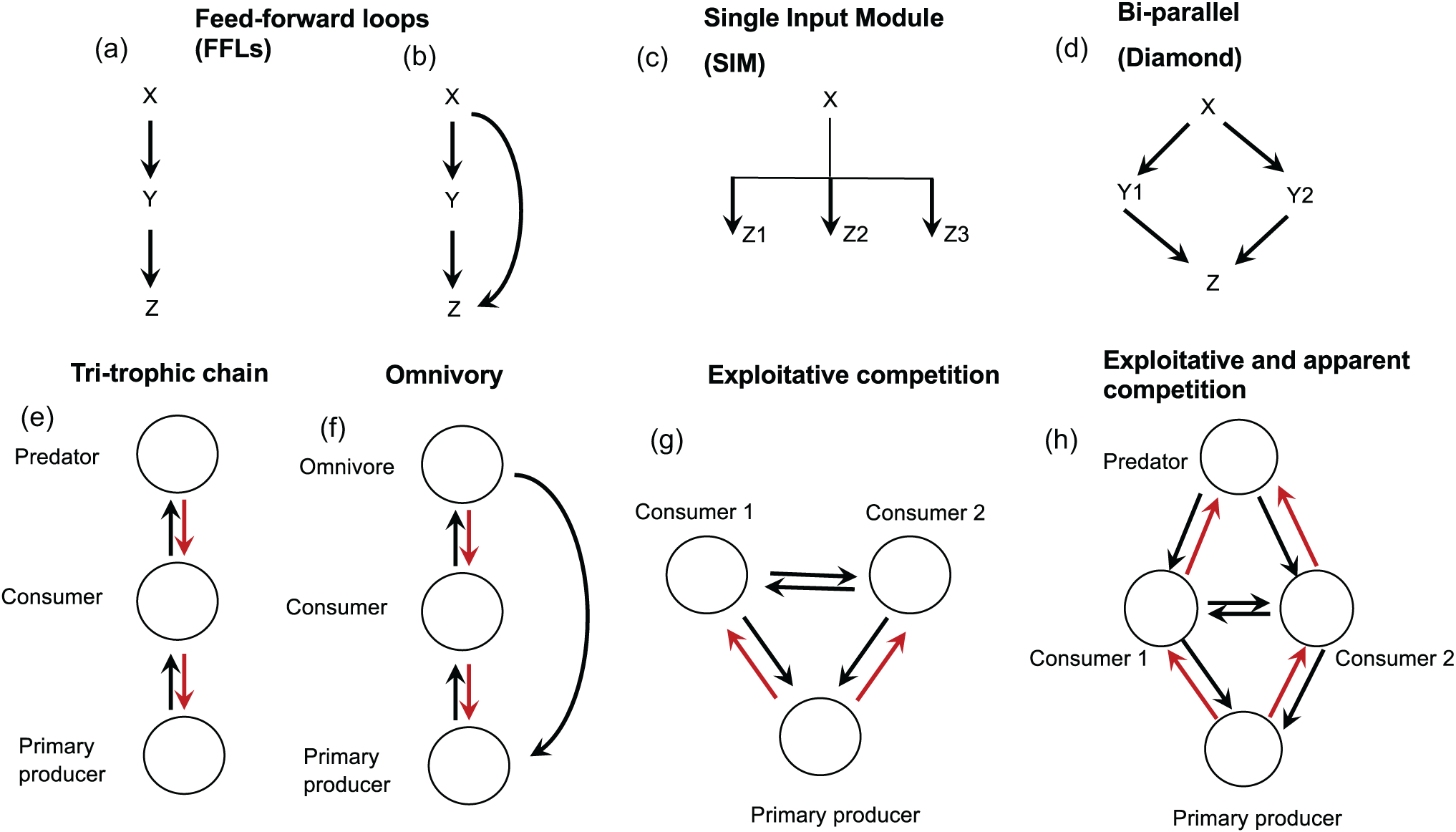
Dominant motifs in biological networks. The top row depicts the Feed-forward loops (FFLs) and Single Input Modules (SIMs) found in transcription networks, and the bi-parallel (diamond) motif found in signal transduction networks. In the FFLs, *X* and *Y* are transcriptional activators and *Z* is a promotor (Alon, 2007); In SIMs, *X* is a regulator and *Z*_*i*_ *i* = 1,, 3 are a group of target genes. In the bi-parallel motif, *X, Y*_*i*_(*i* = 1, 2), *Z* represent signaling proteins and the arrows represent processes such as phosphorylation (Alon, 2007). The bottom row depicts the equivalent food web motifs: try-trophic food chain, omnivory, exploitative competition and the diamond motif arising from the combination of exploitative and apparent competition. Note that the ecological motifs constitute feed-forward loops in terms of energy transfer but feedback loops in terms of interactions among consumers and resources.

A substantial body of theory has been developed to investigate community stability and complexity (e.g., McCann (2000); Williams and Martinez (2000); Rossberg et al. (2005, 2006*b*,*a*); Williams and Martinez (2008); Rossberg et al. (2005, 2006*b*,*a*); McCann (2011); Stouffer and Bascompte (2011)). Nearly all of these involve developing mathematical models to describe the structure and properties of food webs. The resulting predictions are compared with statistical measures of equivalent properties of empirical food webs. One approach generates predator and prey distributions using evolutionary processes of speciation, extinction and migration, which are then compared with equivalent distributions of real food webs (Rossberg et al., 2005, 2006*b,a*). A second approach uses structural models that do not explicitly model species interactions but use mechanistic rules to reproduce food web properties (Williams and Martinez, 2000, 2008; Cattin et al., 2004; Stouffer et al., 2005; Allesina and Pascual, 2008). In these models species are randomly assigned to a niche axis such that the niche values form an ordered set, with each species having a certain probability of feeding on those with lower niche values. This framework includes the niche model (Williams and Martinez, 2000, 2008), the nested hierarchy model (Cattin et al., 2004), the generalized cascade model (Stouffer et al., 2005), and the minimum potential niche model (Allesina and Pascual, 2008). A third approach utilizes random matrix theory to derive stability properties of simple food webs under the assumptions of stable point equilibria and self-limitation in all interacting species (e.g., Allesina and Tang (2015); Monteiro and Faria (2016)). A fourth approach uses explicit dynamical models of species interactions, typically employing the generalized Lotka-Volterra model (e.g., Thébault and Fontaine (2010)) and bioenergetic versions thereof (e.g., McCann (2011); Stouffer and Bascompte (2011)).

As might be expected, given the diversity of approaches, these studies report conflicting findings. Rule-based structural models suggest that omnivory is unlikely to be a stable motif (e.g., Johnson et al. (2014)), while dynamical models suggest it to be strongly stabilizing (e.g., McCann (1997); McCann et al. (1998). Some structural models find apparent competition to be overrepresented (e.g., Bascompte and Melian (2005)), while others find it to be underrepresented (e.g., Camacho et al. (2007)). Random matrix approaches that assume stable point equilibria generated by self-limitation in all species find consumer-resource communities to be more stable than mutualistic communities (e.g., Allesina and Tang (2015)). In contrast, dynamical models that explicitly consider the oscillatory nature of consumer-resource interactions find consumer-resource interactions to be highly unstable (extinction-prone) and that weak horizontal links (e.g., competitive interactions) are required to reduce the oscillatory tendency and increase persistence (e.g., McCann (1997); McCann et al. (1998); McCann (2011)). Even statistical analyses of real communities are divided on the topologies of consumer-resource and mutualistic communities. Most studies find mutualistic communities to have a nested structure (e.g., Bascompte et al. (2003); Bastolla et al. (2009); Suweis et al. (2013) but some do not (e.g., Thébault and Fontaine (2010); Payrató-Borras et al. (2019)). Some studies find consumer-resource communities to be compartmentalized (e.g., Stouffer and Bascompte (2011)), but others find them to be nested (e.g., Kondoh et al. (2010)), or exhibiting a combination of the two topologies (e.g., Kondoh et al. (2010); Thébault and Fontaine (2010)). The diversity of approaches and outcomes has been greatly beneficial in enhancing our understanding of how complex communities can be stable. Further progress, however, requires reconciling these differences to find common ground.

Here we attempt a small first step towards addressing this challenge. We develop a mechanistic framework that combines the topological features of networks with biologically realistic dynamics of consumer-resource interactions. On the topological side we consider trophic interactions as feed-forward loops in which energy is transferred from primary producers to secondary consumers and top predators. On the dynamical side we consider these interactions as feedback loops in which producers have positive effects on consumers, while consumers have negative effects on producers. We use this framework to predict the types of network motifs that emerge under minimal conditions of a constant nutrient input, a single niche axis, no external inputs of nutrients or species, no self-limitation other than that induced by nutrient limitation, and no spatial heterogeneity. Starting from the very basal level — nutrient uptake by primary producers — we investigate which, if any, motifs emerge from the interplay between competition and predation, and whether they are robust to perturbations.

### Conceptual framework

We use theory and data on transcription and signal transduction networks to generate hypotheses about feasible network motifs in ecological communities. As noted above, transcription networks in unicellular organisms (e.g., yeast and *E. coli*; (Alon, 2007)) exhibit feed-forward loops (FFLs) that resemble tri-trophic food chains or a closed loop with omnivory (Fig. 1(a) and (b)). For instance, when the biomass of a primary producer exceeds the level at which a secondary consumer (e.g., herbivore) can persist on it, it opens up the possibility of a primary producer-secondary consumer interaction. Similarly, when primary productivity is sufficiently high to generate a secondary consumer biomass exceeding the level at which a tertiary consumer (e.g., predator) can persist, a primary producer-secondary consumer-tertiary consumer interaction can form, with the primary producer “controlling” the secondary consumer directly, and the tertiary consumer indirectly, through the provision of energy. The persistence of such loops are enhanced when there is negative autoregulation (self-limitation) at one or more trophic levels. When the tertiary consumer can derive energy directly from the primary producer, the latter directly “controls” both consumer trophic levels (Fig. 1). Unicellular transcription networks also exhibit single input modules (SIM), which are akin to exploitative and apparent competition motifs in ecological communities (Fig. 1(c)). When species engage in both exploitative and apparent competition, we get the bi-parallel (diamond) motif found in signal transduction networks (Fig. 1(d)). When two primary producers each support two secondary consumers, we get the bi-fan motif found in neural networks.

Of note, ecological networks are distinct from other biological networks in two ways. First, it is energy, rather than information, that is transferred through the network. Second, species interactions constitute both feed-forward loops and feedback loops. Energy is transferred unidirectionally from primary producers to consumers, with species at lower trophic levels having a positive effect on those at higher trophic levels. At the same time, by extracting energy through direct feeding or other means, species at higher trophic levels have a negative effect on those at lower trophic levels. Similarly, species at the same trophic level compete to acquire energy from the lower trophic level, thus leading to mutually negative effects on one another. It is these feedback loops that define the dynamical nature of consumer-resource communities, with species’ abundances changing as a result of interactions within and between trophic levels.

The next step is to make the connection between motifs and modularity. Networks that are locally cohesive, i.e., the fraction of the feasible edges (links) that occur around a given node (molecules, species), exhibit a high degree of clustering (Watts and Strogatz, 1998). High clustering is a signature of modularity, a group of linked nodes whose collective action achieves a particular function (Milo et al., 2002). In transcription networks, a module is a set of co-regulated genes that share a common function; in signaling pathways, a module is a chain of interacting proteins propagating a signal within a cell (Alon, 2003, 2007). In ecological communities, a module is a group of interacting species whose collective action (energy acquisition) leads to production of biomass.

One could hypothesize that motifs such as FFLs, SIMs and bifans are common in unicellular organisms because they represent the set of minimal modular configurations that can both emerge in a closed system and are robust to perturbations. If this is the case, the preponderance of tritrophic chains, and to some extent omnivory, in ecological communities could be because these motifs constitute the feasible configurations that can both emerge in closed communities and are robust to species invasions. Below we develop this hypothesis in more detail.

Consider a community with a constant nutrient input, in which the total nutrient availability sets the upper limit to the total biomass, and hence the number of species the community can contain. The ways in which species apportion the available biomass determines the number of species and the types of interactions that the community contains. The basal unit is a nutrient-primary producer interaction. In what follows we refer to the primary producer as a plant, but the ideas we develop are general and can apply equally well to other primary producers such as phytoplankton. The plant species’ growth and reproduction depends on an essential nutrient (e.g., Nitrogen, Phosphorous), which it converts into biomass, and for which the individuals in the population compete. The plant will compete with other plant species for a common nutrient pool, and be subject to attack by herbivores. These herbivores in turn are consumed by predators that can also be omnivorous (Fig. 1).

A motif that emerges out of such an interaction has to satisfy two criteria. The first is its feasibility. The possible set of species interactions have to follow the order of nutrient and energy flow, and trophic status. To give an obvious example, we cannot have a consumer without a basal resource, just as we cannot have a top predator without an intermediate consumer. The second criterion is robustness. A stable motif is one that is both (i) stable to perturbations of its constituents’ abundances, and (ii) cannot be replaced by another configuration, i.e., it is robust to species additions or removals. Of note, perturbations may lead to substitutions of species occupying a particular position of the motif, but they will not alter its configuration. For instance, if a tri-trophic food chain is a stable motif, a second herbivore can either invade and exclude the resident herbivore or be excluded by the latter; it cannot invade and coexist with the resident herbivore species. We quantify stability in terms of permanence (long-term persistence of all interacting species), which also encompasses the notion of mathematical stability (return to a non-trivial steady state following a perturbation of species’ abundances).

We make two predictions. First, in a closed community with a constant energy input in which the primary producer’s growth depends on a single limiting nutrient, the emergent motifs are vertical chains (nutrient-plant-herbivore, nutrient-plant-herbivore-predator) with the single exception of a closed loop in the case of omnivory. We expect the frequency of vertical chains to exceed that of omnivory because more ‘coherent’ webs in which species feed on only one trophic level are more stable to perturbations than less coherent ones (Johnson et al., 2014). Second, because vertical chains exhibit high cohesiveness (i.e., the ratio of realized to allowable links approaches 1), they also constitute modules that achieve the common function of biomass production. Modules also tend to localize perturbation impacts, thus increasing the community’s robustness to perturbations. Below we explain the rationale underlying these expectations.

We know from competition theory that, in the absence of local niche partitioning via multiple limiting factors (Tilman, 1982; Chesson, 2000), environmental heterogeneity that allows for spatial or temporal niche partitioning (Chesson, 2000; Amarasekare, 2003), allochthonous nutrient inputs, or dispersal that allows for species recolonizations or source-sink dynamics (Chase and Leibold, 2003; Leibold et al., 2004; Amarasekare, 2003), the species that reduces its resource to the lowest level will exclude all other species (*R*^*\**^; Tilman (1982)). This is a mechanistic interpretation of the competitive exclusion principle that applies to producers and consumers alike. In the absence of ameliorating factors, there can be only as many species at any given trophic level as there are resources at the level below.

The *R*^*\**^ rule means that, in a closed community with a single limiting factor, a second plant species cannot invade and coexist with the resident. Even in the case that the plant species are active at different times and partition the nutrient supply in time, a herbivore entering the community, save in the unlikely event of identical preferences for both plant species, will cause the exclusion of the plant species more susceptible to herbivory (*P* ^*\**^; (Holt, 1977)). Hence, diversity can increase only through the addition of a vertical link to the initial nutrient-plant interaction. This link can be either mutualistic (e.g., pollinator, seed disperser) or antagonistic (e.g., herbivore). Here we focus our attention to antagonistic interactions.

The nutrient-plant community can be invaded by a herbivore if standing plant biomass exceeds that required for the herbivore to maintain itself. The *R*^*\**^ rule ensures that only a single species can occupy the secondary consumer trophic level: the herbivore that reduces the plant biomass to a lower level will exclude all other invaders. Temporal partitioning may allow two herbivores to coexist on the plant, but the arrival of a top predator will exclude the herbivore more susceptible to the predator (*P* ^*\**^ rule). A second vertical link is therefore the most likely configuration in an closed community. Omnivory (i.e., a species feeding on both plant and herbivore trophic levels) can convert the tri-trophic chain into a closed loop (Fig. 1). Below we formalize these predictions mathematically.

### Mathematical model

The dynamics of a consumer-resource community are given by:

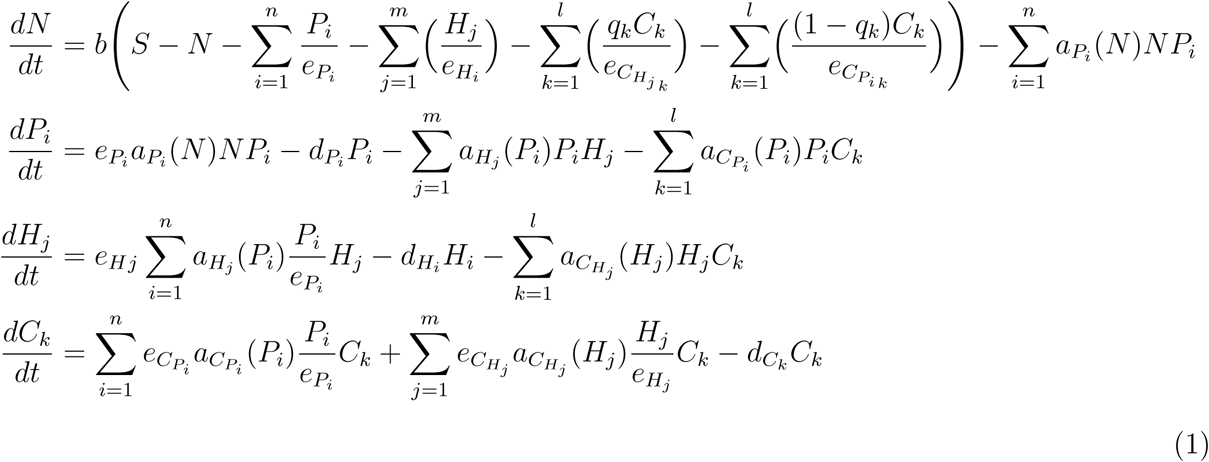

where *S* is the total nutrient content in the system, *b* is the nutrient turnover rate, *N* is the nutrient availability at any given time, and *P*_*i*_, *H*_*j*_ and *C*_*k*_ are, respectively, the biomasses of the *i*^th^ plant species, *j*^th^ herbivore and *k*^th^ predator/omnivore. Since the system is closed, the total nutrient content *S* remains constant over time, imposing a mass balance constraint (Loreau, 1994, 1995) on the system such that

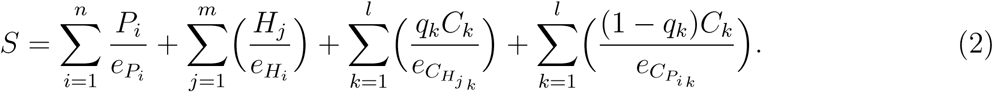

The function *a*_*X*_(*X*) *X* = *P*_*i*_, *H*_*j*_, *C*_*k*_ is the per capita uptake rate, which can be linear (*a*_*X*_(*X*) = *a*_*X*_) or saturating 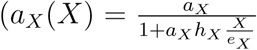 where *h*_*X*_ is the handling time). Importantly, *a*_*X*_(*X*) represents matter and energy flow through the system (Loreau, 1994, 1995). Although the model does not explicitly consider energy, the flow of matter to producers and consumers is dependent on the flow of energy from photosynthesis (Loreau, 1995); the amount of energy available to producers and consumers is, therefore, encapsulated in *a*_*X*_(*X*). The parameters *d*_*X*_(*X* = *P, H, C*) and *e*_*X*_ depict respectively, the per capita mortality rate and the unit (e.g., gram) of biomass generated per unit of nutrient. The fraction 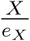 is, then, the total nutrient amount contained in species *X*. Note that *q* is the proportion of the predator’s biomass from feeding on the herbivore, and 1 *− q*, the proportion from feeding on the plant. The magnitude of *q* determines the strength of omnivory.

Equation (1) provides a mechanistic representation of exploitative competition based on the *R*^*\**^ rule (Tilman, 1982). Species at each level (plant, herbivore, predator) compete for a resource whose dynamics are explicitly modelled. For instance, plants compete for nutrients, herbivores compete for plants, predators compete for herbivores, etc. Competition is experienced through the effects that other species have on the abundance of the common “resource”. For brevity, we will refer to the primary producer as plant, and to herbivores and predators as antagonists.

Our approach is three-fold. First, we investigate the emergence of stable motifs via community assembly from the ground up, starting with a plant species that colonizes an empty habitat whose establishment facilitates subsequent invasions by competitors and antagonists. We define a stable motif as one whose configuration, once attained, is robust to species additions and removals. We use mathematical invasion analysis (i.e., the ability of an incoming species to increase from initially small numbers when the resident community is at equilibrium) to quantify robustness. Invasion analyses have the drawback that they focus on the stability of a resident community to a single invader. The mathematical methods involved do not easily lend themselves to investigating the outcomes when more than one species can simultaneously enter a community. In our second approach we use numerical analyses to determine which species combinations persist in the long term when two or more species simultaneously invade a community. If a given motif were truly robust to perturbations, we would expect it to be stable to invasions by single and multiple species. Our third approach to quantifying robustness is species sorting and community disassembly. In the case of sorting, we initiate each community with the full complement of species, allow interactions to proceed, and determine which configurations are persistent. In the case of disassembly, we start with the full complement of species and sequentially remove competitors, antagonists, and primary producers. We determine which motifs remain stable to species removals.

### Model analysis

We use a combination of analytical methods and numerical simulations to investigate community assembly, species sorting, and community disassembly. In the simpler cases of community assembly (e.g., two and three-species interactions), we use mathematical invasion analyses to derive the conditions under which an incoming species can maintain a positive per capita growth rate when the resident species are at equilibrium. Details of these analyses are given in the online Appendix A. We investigate the more complex cases of community assembly and all cases of sorting and disassembly using extensive numerical simulations of the biologically feasible parameter space. In the case of species invasions, we initiate the community at the boundary equilibrium in the absence of the invader(s), and introduce the invader(s) once the resident community has reached its steady state. In all cases the initial invader abundance was set to one individual. In the case of sorting, we initiate each community with the full complement of species (e.g., two plant species, two herbivore species, one predator/omnivore), allow interactions to proceed for 25, 000 time units, and determine which configurations are persistent. In the case of disassembly we allow the full community to interact for 25, 000 time units and, after determining that the community has achieved a steady state, sequentially remove competitors, antagonists, and primary producers. We determine which motifs remain stable to species removals. In all cases, we consider a given interaction to be stable to species additions or removals if its topology (e.g., vertical chain, triangle, diamond) remains intact (i.e., resident species are either resistant to invasion or invading species replace the residents without altering the topology). Parameter definitions and values are given in Table 1.

**Table 1:**
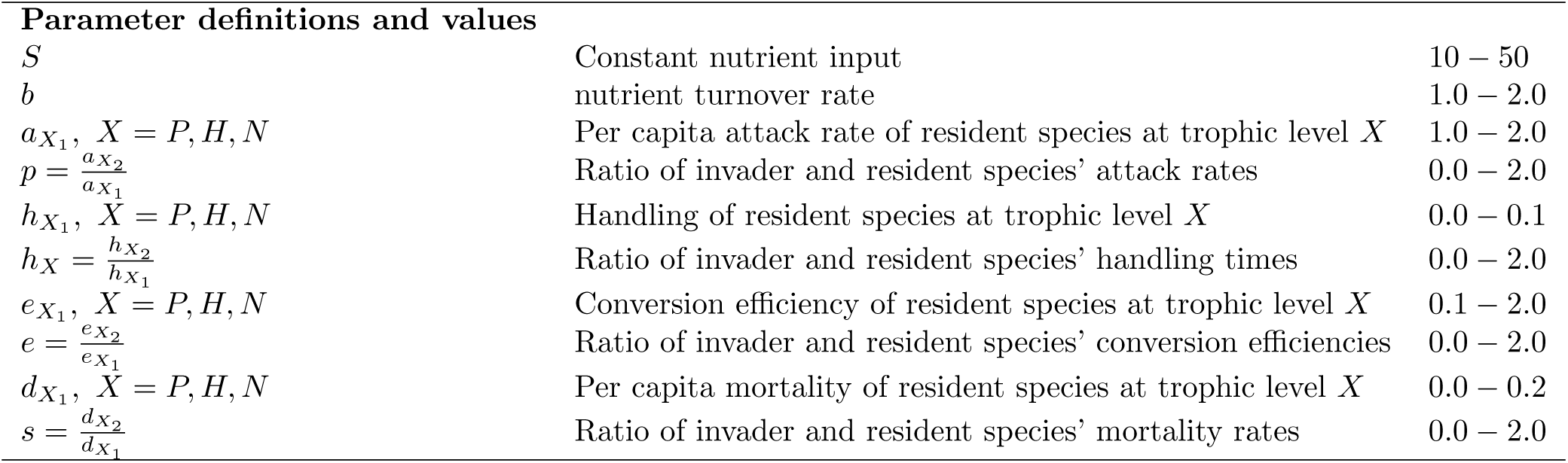
Parameter definitions and values

| Parameter definitions and values |  |  |
| --- | --- | --- |
| $S$ | Constant nutrient input | 10 – 50 |
| $b$ | nutrient turnover rate | 1.0 – 2.0 |
| $a_{X_1}, X = P, H, N$ | Per capita attack rate of resident species at trophic level $X$ | 1.0 – 2.0 |
| $p = \frac{a_{X_2}}{a_{X_1}}$ | Ratio of invader and resident species' attack rates | 0.0 – 2.0 |
| $h_{X_1}, X = P, H, N$ | Handling of resident species at trophic level $X$ | 0.0 – 0.1 |
| $h_X = \frac{h_{X_2}}{h_{X_1}}$ | Ratio of invader and resident species' handling times | 0.0 – 2.0 |
| $e_{X_1}, X = P, H, N$ | Conversion efficiency of resident species at trophic level $X$ | 0.1 – 2.0 |
| $e = \frac{e_{X_2}}{e_{X_1}}$ | Ratio of invader and resident species' conversion efficiencies | 0.0 – 2.0 |
| $d_{X_1}, X = P, H, N$ | Per capita mortality of resident species at trophic level $X$ | 0.0 – 0.2 |
| $s = \frac{d_{X_2}}{d_{X_1}}$ | Ratio of invader and resident species' mortality rates | 0.0 – 2.0 |

## Results

### Community assembly: invasion by one species at a time

1. **Invasion by a plant species** One would expect initial colonizers of empty habitats (e.g., early successional plant species) to exhibit strategies for reproduction and seed dispersal that do not depend on mutualistic partners. Examples involve obligately selfing plant species with wind-dispersed seeds, and facultatively outcrossing species that can revert to selfing in the absence of animal pollinators. In either case, a plant species can increase when rare and reach a steady state with the nutrient as long as the nutrient input exceeds the level to which the plant species suppresses the nutrient at equilibrium, i.e.,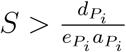 (Appendix A).
2. **Nutrient-plant (NP) community: invasion by a herbivore** A herbivore can successfully invade a nutrient-plant community provided the plant biomass at the steady state with the nutrient exceeds the level to which the herbivore would suppress it, i.e., 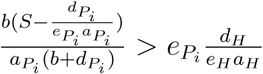 (Fig. 2(a), Appendix A, Fig. S1).
3. **Nutrient-plant-herbivore (NPH) community: invasion by a top predator** A predator can invade a plant-herbivore community provided the herbivore biomass at the nutrient-plant-herbivore steady state exceeds the level to which the predator would depress it (Fig. 2(b), Appendix A, Fig. S1).
4. **Nutrient-plant-herbivore (NPH) community: invasion by an omnivore** An omnivore can invade a plant-herbivore community provided the plant and herbivore biomass at the nutrient-plant-herbivore steady state exceeds the level that the omnivore requires to maintain itself (Fig. 2(c), Appendix A, Fig. S1). However, invasion leads to the exclusion of the herbivore. Coexistence via omnivory is much less frequent (Fig. 2(c)). This is because the omnivore has the advantage of feeding on two trophic levels, while the herbivore has the constraint of feeding only on the level below while being fed on by the level above. Exclusion of the herbivore occurs even when relative non-linearity (Armstrong and McGehee, 1980), mediated via Type II functional responses in the plant, herbivore and omnivore, allows an additional coexistence opportunity (Fig. S2).
5. **Addition of horizontal links** Consistent with expectations, when plant growth is limited by a single essential nutrient, the operation of *R*^*\**^ and *P* ^*\**^ rules prevent the formation of horizontal links even when plant and herbivore species exhibit trade-offs in resource acquisition ability or susceptibility to herbivory. This is because such trade-offs can only increase fitness differences between species (i.e., differences in per capita growth rates in the absence of density-dependent feedbacks; Chesson (2000)); they cannot in themselves generate the stabilizing negative feedbacks that allow species to limit themselves more than they do others (Chesson, 2000). Generating such feedbacks requires more than one niche dimension. In a closed system with a single limiting nutrient and no spatial heterogeneity, the only possible dimension is time. As noted above, temporal partitioning of the basal nutrient by two plant species may generate a horizontal link, but this link is susceptible to invasion by a herbivore, which would set the *P* ^*\**^ rule in motion. To see this consider the two possible mechanisms of temporal partitioning. First, if the two plant species respond differentially to temporal variation such that one species has a high nutrient uptake rate during periods of the year when the other species exhibits little or no activity (e.g., spring and summer annuals), they may coexist via temporal niche partitioning (Fig. 3, Appendix B, Fig. S2). Second, if the plant species differ in the non-linearity of their resource uptake functions such that the species with the more non-linear response generates oscillations in nutrient-plant abundance, a second plant species with a less non-linear response can invade and coexist through the mechanism of relative non-linearity (Armstrong and McGehee, 1980). Coexistence via relative non-linearity can occur when the species with the less non-linear response is better at utilizing the nutrient when it is rare, and the species with the more non-linear response is better at utilizing the nutrient when it is abundant (Fig. 3, Appendix B, Fig. S2). However, neither coexistence mechanism is stable to invasion by a herbivore. Such invasion leads to the exclusion of the plant species more susceptible to herbivore attack (Fig. 3, Fig. S2). As expected, *R*^*\**^ and *P* ^*\**^ rules prevent the formation of horizontal links when the nutrient-plant-herbivore (*NPH*) interaction is invaded by additional plant or herbivore species. The same goes for the tri-trophic chain (*NPHC*) and omnivory (*NPHO*). Below we provide details of each case.
  i. When the *NPH* interaction is invaded by a second plant or herbivore species, two outcomes are possible: the original *NPH* interaction is stable to invasion, or the invading plant or herbivore species replaces the resident species (Fig. 4). There are no species additions at plant or herbivore trophic levels. These results are robust to plant and herbivore species having linear functional responses or saturating ones that allow for relative non-linearity (Figs. 4 and S3).
  ii. The same two outcomes occur when the tri-trophic food chain (*NPHC*) is invaded by a second plant or herbivore species. However, invasion failure is more frequent than replacement of the resident plant and herbivore species by the invaders (Fig. 5).
  iii. In contrast to the linear chains (*NP, NPH, NPHC*), omnivory is not stable to invasions by additional plant or herbivore species (Fig. 5). This is because omnivory itself is rare in closed systems with a limited nutrient supply (see above). Invasion by a second plant species does not alter the omnivore’s advantage of being able to feed on two trophic levels. The herbivore is excluded, and the plant species that is less susceptible to omnivore attack will exclude the other. The overall outcome is a nutrient-plant-omnivore interaction with the omnivore relying solely on herbivory. Similarly, invasion by a second herbivore species results in the exclusion of the inferior competitor for the common plant resource (Fig. 5).
6. **Community assembly via invasion by single species: summary of results** When community assembly occurs in the absence of niche partitioning mechanisms that allow the addition of horizontal links, increase in diversity can occur only through the addition of a vertical link. If the plant species’ per capita growth rate does not depend on a mutualist (e.g., because it is obligately selfing or has wind-dispersed seeds), the first vertical link is most likely be a herbivore, which opens up the possibility of invasion by a top predator or an omnivore. Vertical chains (*NP, NPH, NPHC*) are more robust to species invasions than omnivory. This is because omnivory involves resource partitioning, the opportunity for which is constrained in a closed system with a limiting nutrient supply that sets the upper limit to community biomass.

**Figure 2:**
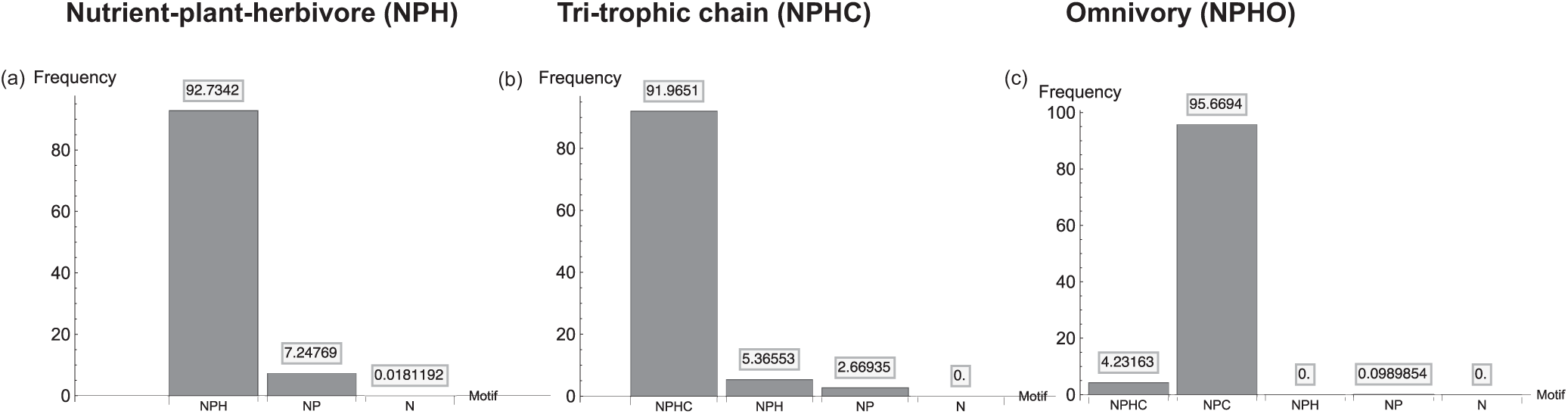
Emergence of vertical chains through sequential species invasions of a nutrient-plant interaction. Panels (a)-(c) depict the frequency distribution of emergent motifs when a nutrient-plant interaction is invaded by a herbivore (panel (a)), a nutrient-plant-herbivore interaction is invaded by a top predator (panel (b)) and nutrient-plant-herbivore interaction is invaded by an omnivore (panels (c)). Parameter definitions and values are given in Tables 1 and 2.

**Table 2:**
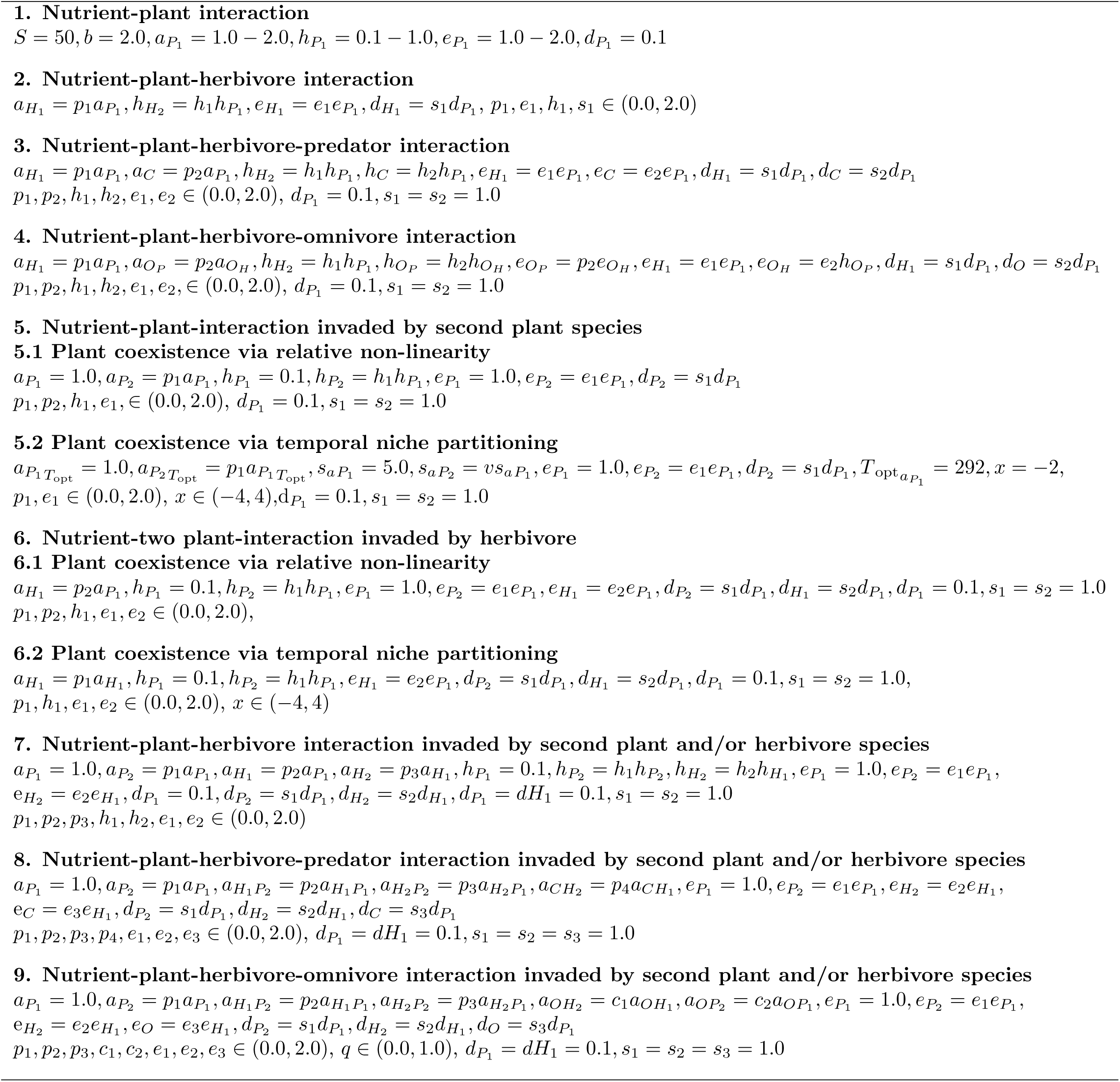
Parameter values used in simulations

**Figure 3:**
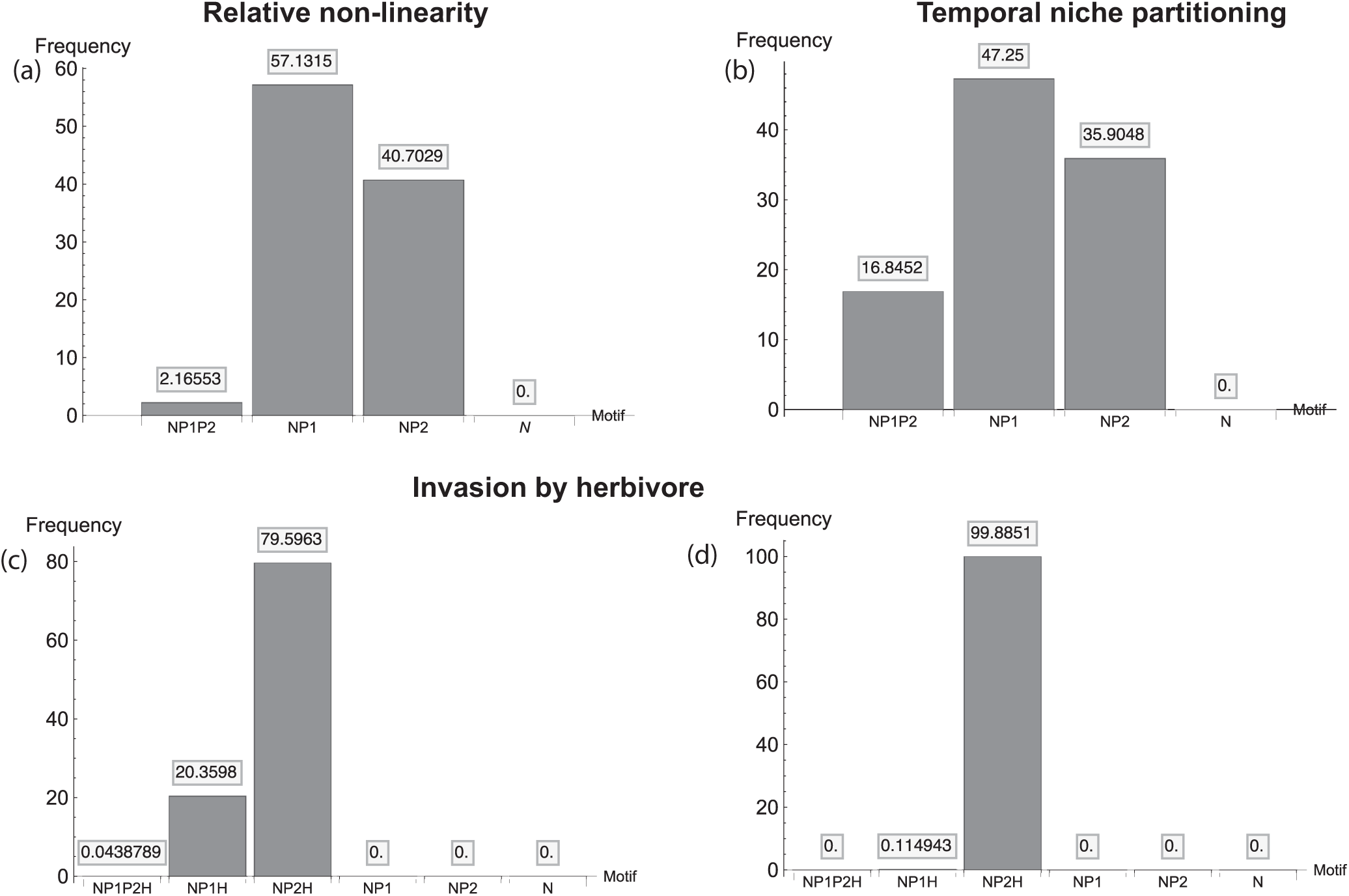
Plant species coexistence via relative non-linearity (RN; panel (a)) and temporal niche partitioning (TNP; panel (b)), and robustness coexistence to invasion by a herbivore (RN: panel (c); TNP: panels (f) and (d)). Parameter definitions and values are given in Tables 1 and 2.

**Figure 4:**
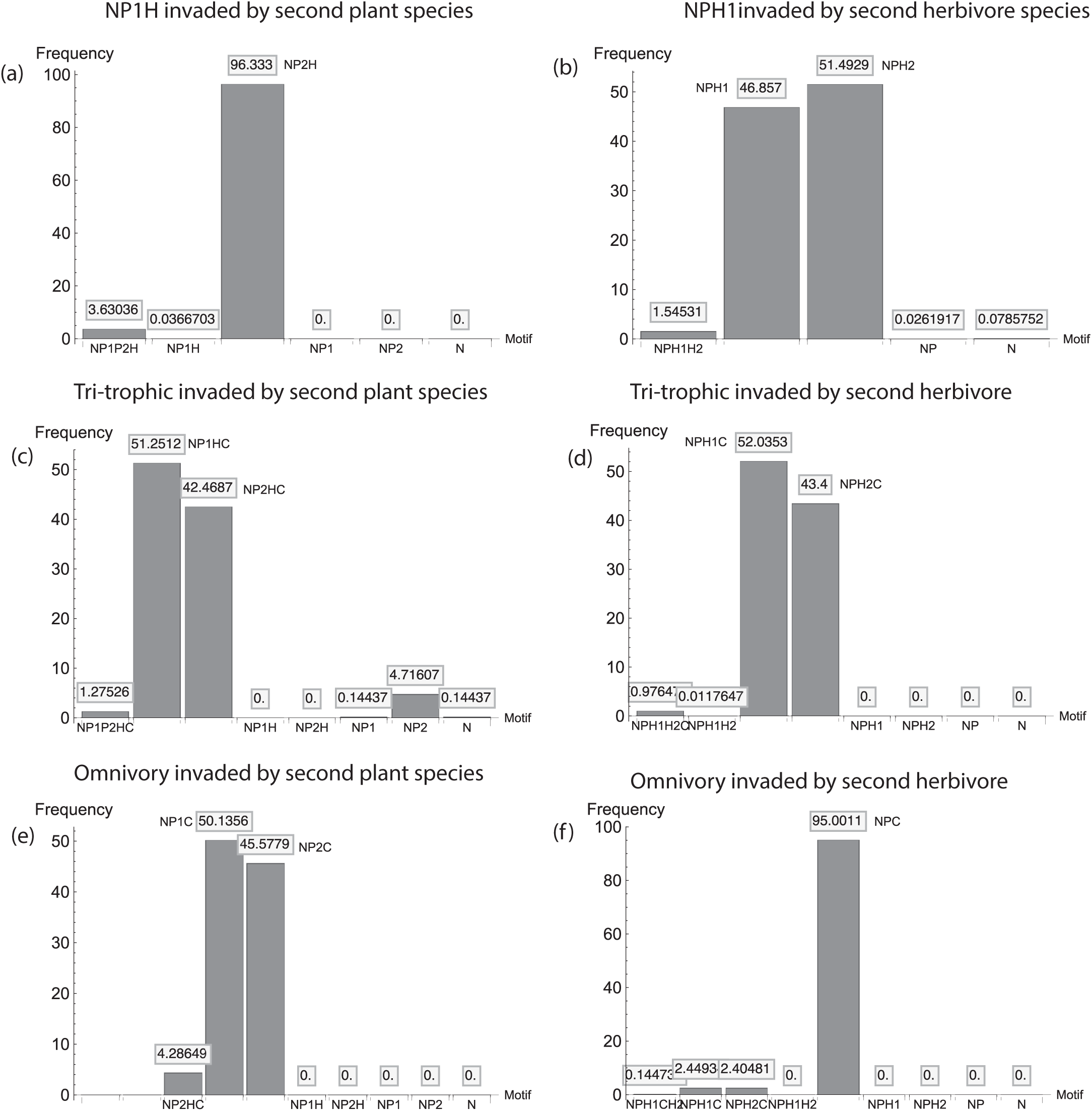
Vertical trophic chains invaded by a second plant or herbivore species. Panels (a) and (b) depict, respectively, the invasion of a NPH interaction by a second plant and second herbivore species. Panels (c) and (d) and (e) and (f) depict the same for the tri-trophic chain and omnivory. Parameter definitions and values are given in Tables 1 and 2.

**Figure 5:**
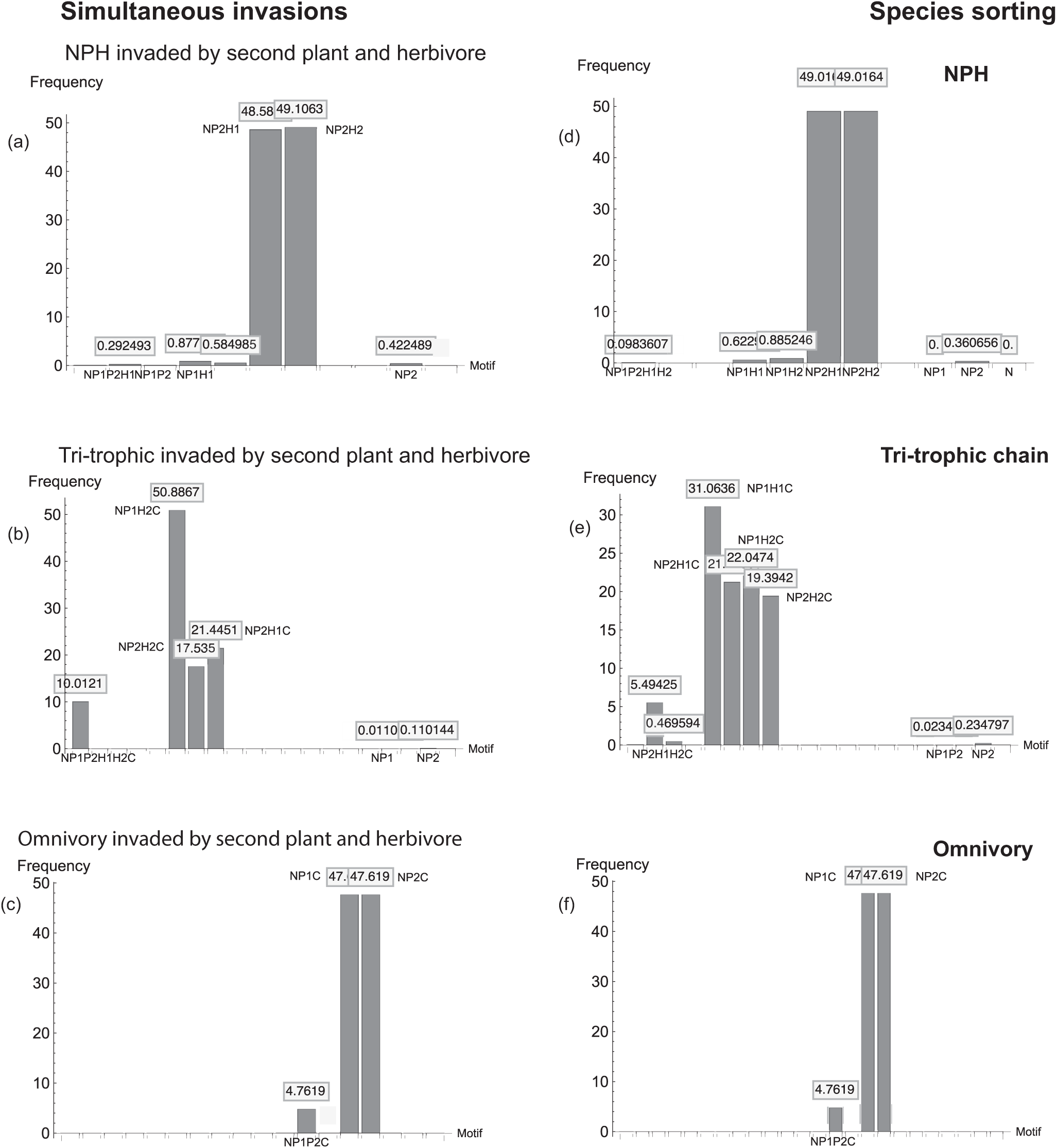
Simultaneous invasion by multiple species and species sorting in vertical trophic chains. Panels in the left column ((a)-(c)) depict simultaneous invasion by a second plant and herbivore species in nutrient-plant-herbivore, tri-trophic, and omnivory interactions. Panels in the right column depict the outcome of species sorting. In the case of sorting, each community is started with the full complement of species (*NP*_1_*P*_2_*H*_1_*H*_2_, *NP*_1_*P*_2_*H*_1_*H*_2_*C, NP*_1_*P*_2_*H*_1_*H*_2_*O*), and allowed to interact for 50, 000 time units. All panels depict the frequency distributions of species in the three communities at the long-term steady state. Parameter definitions and values are given in Tables 1 and 2.

### Community assembly: simultaneous invasion by multiple species

As noted previously, mathematical invasion analysis focuses on the conditions under which a single species can increase from initially small numbers when the rest of the community is at a steady state. In reality, more than one species can enter a community at any given time. Numerical simulations spanning a large parameter space show that vertical chains (*NPH, NPHC*) are robust to the simultaneous invasion of competitors and antagonists, but omnivory (*NPHO*) is not (Fig. 5). Below we explain these results in detail.

1. **Nutrient-plant-herbivore (***NPH***) community: invasion by plant and herbivore species** When plant and herbivore species simultaneously invade the *NPH* community, the outcome is invasion failure or the replacement of resident species by invaders (Fig. 5). This is a direct result of the combined operation of *R*^*\**^ and *P* ^*\**^ rules. The plant species that can, overall, extract more of the nutrient supply while evading or resisting herbivory will exclude the other. In the case of the herbivore, the species that reduces plant biomass to the lowest level will exclude the other. While it is possible for one plant species to be the superior nutrient competitor and be more susceptible to herbivory, such trade-offs generate only fitness differences and not the the stabilizing niche partitioning mechanisms required for coexistence (Chesson, 2000).
2. **Nutrient-plant-herbivore-predator (***NPHC***) community: invasion by plant and herbivore species** The outcome is the same as that for the NPH community. The plant species that extracts more of the nutrient in the face of herbivory will exclude the other, and the herbivore that can consume as much plant biomass as possibly while evading predation will exclude the other (Fig. 5). Invasion by a second top predator in combination with a plant or herbivore species leads to the same outcome.
3. **Nutrient-plant-herbivore-omnivore (***NPHO***) community: invasion by plant and herbivore species** In contrast to the vertical chains, omnivory proves to be unstable to simultaneous invasion by multiple species. Regardless of which combination of species invades (plant and herbivore, plant and omnivore, herbivore and omnivore), the outcome is the exclusion of the herbivore and the emergence of a vertical chain with the omnivore acting as a top predator (Fig. 5). This is because, as shown above in the single invasion case, the omnivore has the advantage of feeding on multiple trophic levels while the herbivore has the dual disadvantage of competing with the omnivore for the plant resource while also being fed on by omnivore. In a closed system with a limiting nutrient supply and no external inputs, the herbivore being a superior competitor for the plant species does not generate sufficient niche partitioning opportunities to allow for herbivore-omnivore coexistence.

### Species sorting

In this step we start with the full assemblage of species for each of the three communities (*NPH, NPHC* and *NPHO*) and allow the dynamics to proceed naturally. We find that species sorting occurs via the operation of *R*^*\**^ and *P* ^*\**^ rules, with the result that the stable motifs to emerge are, again, the vertical chains (*NPH, NPHC* and *NPO*).

1. **Nutrient-two plant-two herbivore community (***NP*_1_*P*_2_*H*_1_*H*_2_) Starting from the full community, species sorting leads to the emergence of the *NPH* chain (Fig. 5). Which plant species persists depends on the cumulative effect of resource acquisition ability and susceptibility to herbivory. Which herbivore species persists depends on the level to which each herbivore can depress plant biomass.
2. **Tri-trophic chain (***NP*_1_*P*_2_*H*_1_*H*_2_*C*) Starting from the full community, species sorting leads to the emergence of the *NPHC* chain as the dominant motif (Fig. 5). Other motifs that occur in low frequency (e.g., *NP*_1_*P*_2_, *NP*_*i*_*H*_1_*H*_2_, *i* = 1, 2) are the result of transient coexistence of species, driven by strong trade-offs, at plant and herbivore trophic levels.
3. **Omnivory (***NP*_1_*P*_2_*H*_1_*H*_2_*O*) As with assembly, species sorting leads to the exclusion of the herbivore by the omnivore, with the vertical chain (*NPO*) being the emergent outcome (Fig. 5).

### Community disassembly

Here we start with the full complement of species for each community, and sequentially remove species starting with the highest trophic level. We find that, across all community types, the motifs that are robust to disassembly are the vertical chains (*NPH, NPHC* and *NPO*). When fitness differences between species are strong, transient coexistence of plant or herbivore species can occur, but this outcome is restricted to the nutrient-plant-herbivore community (*NPH*); it is not observed in *NPHC* or *NPHO* interactions.

1. **Nutrient-two plant-two herbivore community (***NP*_1_*P*_2_*H*_1_*H*_2_) When one herbivore species is removed, the community simplifies to two nutrient-plant-herbivore chains (*NP*_*i*_*H, i* = 1, 2) (Fig. 6(a) and (b)). Removal of one plant species also leads to the formation of nutrient-plant-herbivore chains (*NPH*_*i*_ *i* = 1, 2; Fig. 6(c) and (d)). Simultaneous removal of one plant and one herbivore species leads to the same outcome (Fig. 6(e) and (f).)
2. **Tri-trophic chain (***NP*_1_*P*_2_*H*_1_*H*_2_*C*) When one herbivore species is removed, the community simplifies to two nutrient-plant-herbivore chains (*NP*_*i*_*H, i −* 1, 2) (Fig. 6(a) and (b)). Removal of one plant species also leads to the formation of nutrient-plant-herbivore chains (*NPH*_*i*_ *i* = 1, 2; Fig. 6(c) and (d)). Simultaneous removal of one plant and one herbivore species leads to the same outcome (Fig. 6(e) and (f)).
3. **Omnivory (***NP*_1_*P*_2_*H*_1_*H*_2_*O*) When one herbivore species is removed, the community simplifies to two nutrient-plant-omnivore chains (*NP*_*i*_*O i* = 1, 2) with the omnivore excluding the remaining herbivore (Fig. 7(a) and (b)). When one plant species is removed, we get a single nutrient-plant-omnivore chain (*NPO*; Fig. 7(c) and (d)). Simultaneous removal of one plant and one herbivore species leads to the same outcome (Fig. 7(e) and (f)). When the omnivore itself is removed, the community simplifies to one of four nutrient-plant-herbivore chains (*NP*_*i*_*H*_*j*_ *i, j* = 1, 2).

**Figure 6:**
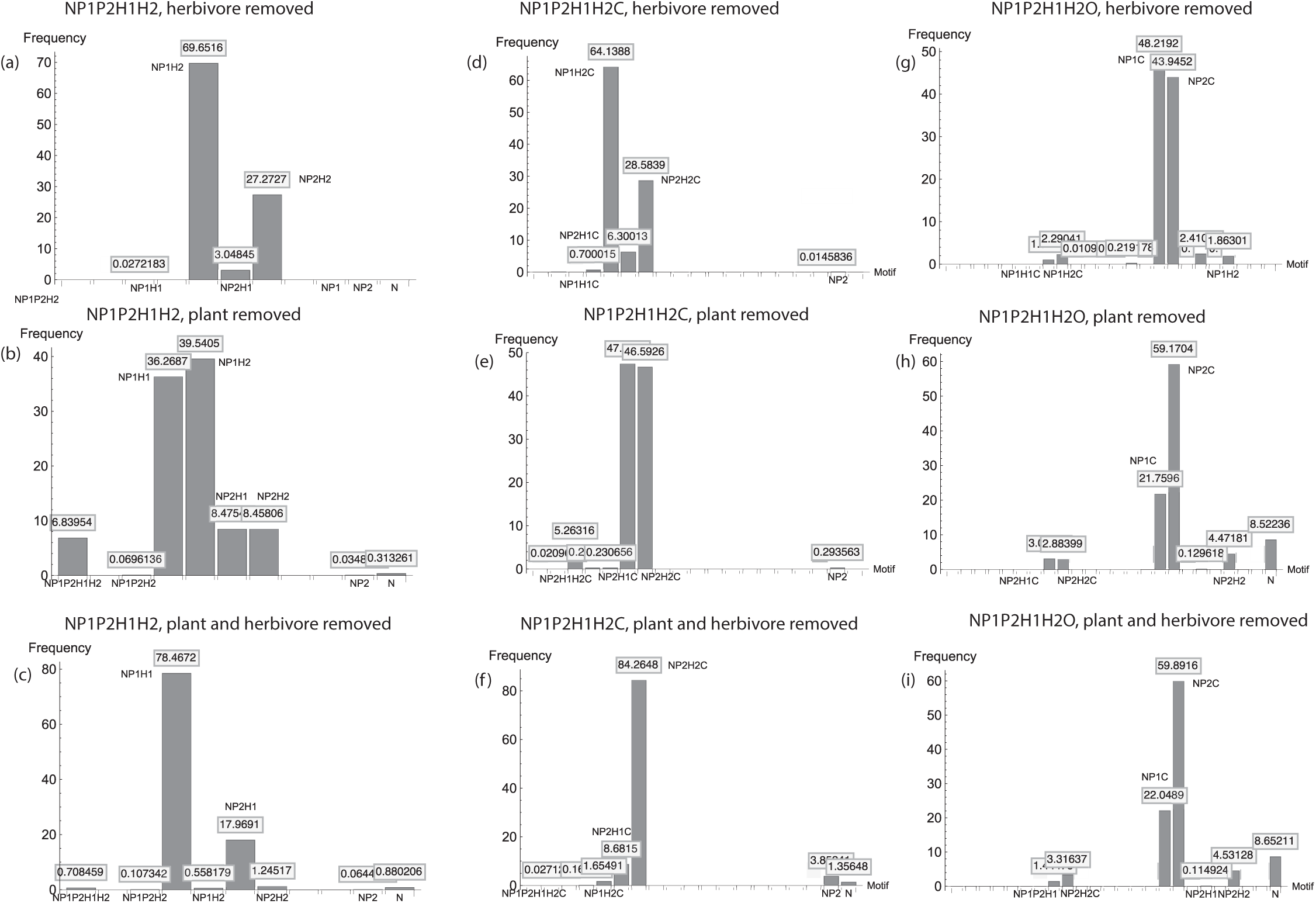
Disassembly of nutrient-plant-herbivore (panels (a)-(c)), tri-trophic (panels (d)-(f)), and omnivory ((g)-(j)) communities. Each community is started with the full complement of species (*NP*_1_*P*_2_*H*_1_*H*_2_, *NP*_1_*P*_2_*H*_1_*H*_2_*C, NP*_1_*P*_2_*H*_1_*H*_2_*O*), allowed to reach steady state, and subjected to sequential species removals (omnivore, predator, herbivore, plant). Parameter definitions and values are given in Tables 1 and 2.

### Summary of results

Taken together, the outcomes of community assembly, species sorting, and community disassembly show that in a closed system with a constant supply of a limiting nutrient, no spatial heterogeneity, and no external nutrient inputs or immigration, the only motifs that can emerge are vertical chains of nutrient-plant-herbivore or nutrient-plant-herbivore-predator. This is true even in the presence of temporal niche partitioning at the plant level and relative non-linearity at plant and herbivore trophic levels.

## Discussion

There is strong empirical evidence that ecological communities exhibit recurrent elements (motifs) that characterize other types of biological networks (e.g., transcription, neural signal transduction; Milo et al. (2002); Alon (2003); Bascompte and Melian (2005); Kashtan and Alon (2005); Alon (2007)). Analyses of network properties in a wide variety of food webs have identified four dominant motifs: exploitative competition, apparent competition, omnivory and tri-trophic chain (Milo et al., 2002; Bascompte and Melian, 2005; Camacho et al., 2007). Nearly all studies identify tri-trophic chains as a dominant motif, but there is some disagreement as to whether omnivory is one (Bascompte and Melian, 2005; Camacho et al., 2007; Johnson et al., 2014). A few studies find exploitative and apparent competition to be dominant (Bascompte and Melian, 2005), but the consensus appears to be that they are less prevalent across different types of food webs than tri-trophic chains or omnivory. At least one study (Milo et al., 2002) has found the diamond motif that arises from a combination of exploitative and apparent competition.

In unicellular organisms such as yeast and *E. coli*, transcription, neural and transduction networks exhibit dominant motifs that resemble those seen in ecological communities. For instance, feed-forward loops (FFLs) in transcription networks include motifs resembling tri-trophic chains and omnivory, and single input modules (SIM) that resemble exploitative and apparent competition. Signal transduction networks exhibit a bi-parallel (diamond) motif, akin to the diamond food web arising from the combination of exploitative and apparent competition. Theory suggests that these motifs are dominant because they constitute the optimal configurations for information transfer that are also robust to perturbations (Milo et al., 2002; Kashtan and Alon, 2005; Alon, 2007). Extending this idea to ecological communities in which the transfer is of energy rather than information, we can hypothesize that the dominant motifs observed in ecological communities are those that represent configurations that are both feasible and stable: they can emerge under minimal conditions in closed communities, and they are robust to invasions by other species.

Here we test this hypothesis using a mechanistic framework that combines the topological features of networks with biologically realistic dynamics of consumer-resource interactions. On the topological side we consider trophic interactions as feed-forward loops in which energy is transferred from primary producers to secondary consumers and top predators. On the dynamical side we consider these interactions as feedback loops in which producers have positive effects on consumers, while consumers have negative effects on producers. We use three approaches — community assembly, species sorting, community disassembly — to investigate the robustness of the five motifs (exploitative competition, apparent competition, omnivory, tri-trophic chain and diamond) found to be prevalent in real food webs (Milo et al., 2002; Bascompte and Melian, 2005; Camacho et al., 2007; Johnson et al., 2014; Monteiro and Faria, 2016).

Our work differs from previous studies in several ways. First, instead of assembling communities by randomly assigning species to a niche axis under prescribed rules, we explicitly investigate community assembly from the ground up, starting with a single primary producer (plant) that colonizes an empty habitat. Second, we investigate how, in a nutrient-limited system subject to a mass balance constraint, sequential colonizations by primary and secondary consumers (herbivores and predators) lead to community assembly in the absence of any self-limitation mechanism save that arising from the constant nutrient input. Third, we do not restrict our investigations to situations in which only stable point equilibria are possible. By incorporating non-linear (saturating) functional responses at all trophic levels, we explicitly allow for the oscillatory dynamics that characterize consumer resource interactions. To our knowledge, this is the first time that these biological realities have been incorporated into investigations of food web assembly.

We find that, in a closed community in which the total nutrient availability sets the upper limit to the total biomass, the only motifs that can emerge are vertical chains of nutrient-plant, nutrient-plant-herbivore, and nutrient-plant-herbivore-predator interactions. Although temporal variation, either through relative non-linearity in functional responses or temporal nutrient partitioning can allow the coexistence of plant species, such coexistence is not robust to invasion by a herbivore. The plant species whose susceptibility to herbivory, when averaged over the year (in temporal niche portioning) or nutrient cycle (in relative non-linearity) is lower will exclude the other plant species. The same process happens at the herbivore level. In the absence of a top predator, the herbivore species that depresses plant biomass to the lowest level will exclude all other species; in the presence of a top predator, the herbivore, that can extract more energy from the plant while withstanding predation will exclude all other species.

On the face of it, this result may seem trivial. After all, what we are seeing is the operation of the *R*^*\**^ and *P* ^*\**^ rules. However, looking beneath the surface reveals several important insights. First, just as feed-forward loops maximize information transfer in transcription networks, vertical chains maximize energy transfer in ecological communities. Since the total amount of energy is constant in a closed community, coexistence at any trophic level means the apportionment of the same amount of energy amongst species (since the amount of energy at that level cannot exceed the amount procured by the species best at resource acquisition) with extra losses during conversion of energy to biomass (determined by the conversion efficiency parameter) and mortality. In a vertical chain, more energy is transferred from one level to another because having a single species at each level minimizes energy loss due to biomass conversion and mortality. Thus, a vertical chain also leads to greater overall biomass and hence productivity of the community. The key point is that the ecological constraint imposed by the *R*^*\**^ and *P* ^*\**^ rules not only serves to make the linear chains more robust to perturbations, but they also maximize energy transfer.

The second insight is that the operation of the *R*^*\**^ and *P* ^*\**^ rules increases the trophic coherence of the community. Coherence is determined by the number of trophic levels that a given consumer occupies (Johnson et al., 2014); a top predator feeds only on the trophic level below it (herbivore) but an omnivore feeds on two trophic levels below it (herbivore and plant). The fewer trophic levels a given consumer extracts energy from, the more coherent a network is, and the less self-regulation required to stabilize it (Johnson et al., 2014). This is why we observe the vertical chains to be not only persistent but also stable to perturbations of species’ abundances. Vertical trophic chains not only maximize energy transfer and biomass production, but they are also stable both in the ecological sense (long-term persistence) and the mathematical sense (recovery from perturbations to species’ abundances). What is notable is that stability is achieved through the minimum possible level of self-regulation: a single negative feedback loop at the primary producer level.

Importantly, the vertical trophic chains that emerge as stable motifs also confer modularity. Vertical chains exhibit high clustering coefficients (i.e., the ratio of realized to allowable links approaches 1; Watts and Strogatz (1998)). Clustering is a signature of modularity, a group of a linked nodes with strong interactions (Alon, 2003). Vertical trophic chains satisfy these criteria. They are highly cohesive (i.e., all allowable links are realized), and they achieve the common function of converting energy into biomass. This conversion is what ultimately constitutes community productivity, be they phytoplankton or forests. Because they can convert energy into biomass in the absence of other interactions, and because perturbations occurring at one or more nodes (species) are contained within the chain and not transmitted horizontally, vertical chains constitute the minimal modular structure that can arise even under the restrictive conditions of a limited energy supply that places an upper limit on biomass production.

These results beg the following question: real communities constitute complex webs of interacting species. While they do contain vertical chains, they also contain a multitude of horizontal links. If it is the vertical chains that maximize energy transfer and are robust to perturbations, how do complex communities with numerous horizontal links persist in the face of abiotic (i.e., nutrient influxes, toxins and pollutants) and biotic (invasions of competitors and antagonists) perturbations? We propose the following explanation: vertical trophic chains constitute the dominant motif because they comprise the backbone of all consumer-resource communities. The reason for this is simple. Vertical chains constitute the set of minimal configurations that can assemble under the most restrictive of conditions: a single limiting nutrient, no external energy inputs, no immigration, no spatial heterogeneity, and minimal self-limitation. Vertical chains are what is left when everything is taken away. They are the foundation on which complex structures are built.

It is noteworthy that our theoretical finding is consistent with the empirical finding that tri-trophic chains are the dominant motif in a wide variety of food webs (e.g., Milo et al. (2002); Camacho et al. (2007)). Previous theory has demonstrated that the dynamical behavior of tritrophic chains can be greatly altered when embedded in complex webs (Otto et al., 2007; Kondoh, 2008; Cohen et al., 2010), raising the issue of whether it is possible to identify tri-trophic chains that can persist in the absence of other interactions. Our theoretical finding that vertical trophic chains can persist independently even under the most restrictive conditions lends credence to previous empirical findings that tri-trophic chains are the dominant motif in real food webs.

It is also noteworthy that our findings contradict the empirical findings that omnivory, exploitative and apparent competition, and the diamond web can be common in natural communities. These discrepancies serve to illustrate important biological realties. Consider first, the omnivory motif. There is disagreement in the empirical literature as to whether omnivory constitutes a dominant motif in food webs. Some studies find it to be prevalent while other do not (Schneider et al., 2012; Brose et al., 2019). There is a considerable body of theory showing that a trade-off between resource acquisition ability and susceptibility to omnivory is insufficient to guarantee herbivore-omnivore coexistence (S. and Feissel, 2000; Amarasekare, 2007, 2008). Our findings serve to demonstrate this result in a closed system that is nutrient-limited and subject to a mass-balance constraint. (Interestingly, when omnivory is prevalent, as in size-structured communities (Schneider et al., 2012; Brose et al., 2019), it is because the link between the basal resource and the omnivore is weak.) The inadequacy of fitness differences to provide stabilizing negative feedbacks is also why we do not observe exploitative and apparent competition. The minimalist scenario we explore provides no opportunities for the niche partitioning mechanisms required to sustain such interactions. This is particularly interesting because we do not observe coexistence above the primary producer (plant) level even in the presence of temporal coexistence mechanisms such as relative non-linearity and temporal niche partitioning. When multiple species occupy multiple trophic levels, we need more than two niche dimensions; resource/natural enemy and time are no longer sufficient. We need to invoke space, external resource inputs, and immigration.

The need to move beyond the minimal conditions of limited energy takes us to considerations of the conditions required for the assembly and persistence of complex ecological communities. We propose that, just in the way that macromolecules (e.g., DNA, protein) are formed by the intertwining of molecular chains that subsequently fold into complicated structures held together by relative fragile bonds, complex communities are formed by the coming together of vertical chains that are then held together by relatively fragile horizontal links that can break when the energy inputs that make them possible are removed, or when perturbations to the existing structure occur in terms of species additions or removals. Let us consider a simple example, the coming together of two trophic chains, each supported at the base by a different nutrient or a different supply of the same nutrient separated in space (e.g., soil space occupied by the root systems of individual plants). Since each chain has an independent base (energy input), the *R*^*\**^ and *P* ^*\**^ rules no longer hold, resulting in coexistence at primary producer and secondary consumer levels. A top predator that attacks secondary consumers of both chains, or an omnivore that feeds on a plant species from one chain and a herbivore from the other, will be able to do so without species losses at lower trophic levels, leading to a compartment that now contains multiple motifs: tri-trophic chain, omnivory, exploitative and apparent competition, and the diamond web. Extending our framework to incorporate multiple vertical chains into models with energy limitation and mass balance constraints is an important next step.

## Acknowledgements

This research was supported by the National Science Foundation grant DEB-1949796 to P.A.

## References

Allesina, S., and M. Pascual. 2008. Network structure, predator-prey modules, and stability in large food webs. Theoretical Ecology 1:55–64.

Allesina, S., and S. Tang. 2015. The stability–complexity relationship at age 40:a random matrix perspective. Population Ecology 57:63–75.

Alon, U. 2003. Biological networks: The tinkerer as an engineer. Science 301:1866–1867.

Alon, U. 2007. Network motifs: theory and experimental approaches. Nature Genetics 8:450–461.

Amarasekare, P. 2003. Diversity-stability relationships in multi-trophic systems: an empirical exploration. Journal of Animal Ecology 72:713–724.

Amarasekare, P. 2007. Trade-offs, temporal variation and species coexistence in communities with inrraguild predation. Ecology 88:2720–2728.

Amarasekare, P. 2008. The coexistence of intraguild predators and prey in resource-rich environments. Ecology 89:2786–2797.

Armstrong, R., and R. McGehee. 1980. Competitive exclusion. American Naturalist 115:151–170.

Bascompte, J., P. Jordano, C. Melian, and J. Olesen. 2003. The nested assembly of plant– animal mutualistic networks. Proceedings of the National Academy of Sciences of the USA 100:9383–9387.

Bascompte, J., and C. J. Melian. 2005. Simple trophic modules for complex food webs. Ecology 86:2868–2873.

Bastolla, U., M. Fortuna, A. Pascual-Garcia, A. Ferrera, B. Luque, and J. Bascompte. 2009. The architecture of mutualistic networks minimizes competition and increases biodiversity. Nature 458:1018–1021.

Brose, U., P. Archambault, B. A.D., L. Bersier, T. Boy, J. Canning-Clode, E. Conti, M. Dias, and C. Digel. 2019. Predator traits determine food-web architecture across ecosystems. Nature Ecology and Evolution 3:919–927.

Camacho, J., D. Stouffer, and L. Amaral. 2007. Quantitative analysis of the local structure of food webs. Journal of Theoretical Biology 246:260–268.

Cattin, M., L. Bersier, C. Banasek-Richter, R. Baltensperger, and J. Gabriel. 2004. Phylogenetic constraints and adaptation explain food-web structure. Nature 427:835–839.

Chase, J., and M. A. Leibold. 2003. Ecological niches: linking classical and contemporary approaches. University of Chicago Press.

Chesson, P. 2000. Mechanisms of maintenance of species diversity. Annual Review of Ecology and Systematics 31:343–366.

Cohen, J., D. Schittler, D. Raffaelli, and D. Reuman. 2010. Food webs are more than the sum of their tritrophic parts. Proceedings of the National Academy of Sciences of the USA 106:22335–22340.

Holt, R. 1977. Predation, apparent competition, and the structure of prey communities. Theoretical Population Biology 12:197–229.

Johnson, S., V. Domínguez-García, D. L., and M. Muñoz. 2014. Trophic coherence determines food-web stability. Proceedings of the National Academy of Sciences of the USA 111:17923–17928.

Kashtan, N., and U. Alon. 2005. Spontaneous evolution of modularity and network motifs. Proceedings of the National Academy of Sciences of the USA 102:13773–13778.

Kondoh, M. 2008. Building trophic modules into a persistent food web. PNAS 105:16631–16635.

Kondoh, M., S. Kato, and Y. Sakato. 2010. Food webs are built up with nested subwebs. Ecology 91:3123–3130.

Leibold, M. A., M. Hoyoak, N. Mouquet, P. Amarasekare, J. Chase, M. Hoopes, R. Holt, J. Shurin, R. Law, D. Tilman, M. Loreau, and A. Gonzalez. 2004. The metacommunity concept: a framework for multi-scale community ecology. Ecology Letters 7:601–613.

Lipson, H., J. Pollack, and P. Suh. 2002. On the origin of modular variation. Evolution 56:1549–1556.

Loreau, M. 1994. Mass and energy flow in closed ecosystems: do ecological or mathematical constraints prevail? Journal of Theoretical Biology 168:237–243.

Loreau, M. 1995. Consumers as maximizers of matter and energy flow in ecosystems. American Naturalist 145:22–42.

McCann, K. 2000. The diversity stability debate. Nature 405:228–233.

McCann, K., A. Hastings, and G. R. Huxel. 1998. Weak trophic interactions and the balance of nature. Nature 395:794–798.

McCann, K. S. 1997. Re–evaluating the omnivory–stability relationship in food webs. Proceedings of the Royal Society of London. Series B: Biological Sciences 264:1249–1254.

McCann, K. S. 2011. Food Webs. Princeton University Press.

Milo, R., S. Shen-Orr, S. Itzkovitz, N. Kashtan, D. Chklovskii, and U. Alon. 2002. Network motifs: Simple building blocks of complex networks. Science 298:824–827.

Monteiro, A., and L. Faria. 2016. The interplay between population stability and food-web topology predicts the occurrence of motifs in complex food-webs. Jounal of Theoretical Biology 409:165–171.

Otto, S., B. Rall, and U. Brose. 2007. Allometric degree distributions facilitate food-web stability. Nature 450:1226–1230.

Payrató-Borras, C., L. Hernández, and Y. Moreno. 2019. Breaking the spell of nestedness: The entropic origin of nestedness in mutualistic systems. Physical Review X 9:2160–3308.

Rossberg, A., H. Matsuda, T. Amemiya, and K. Itoh. 2005. An explanatory model for foodweb structure and evolution. Ecological Complexity 2.

Rossberg, A. 2006a. Foodwebsl: experts consuming families of experts. Journal of Theoretical Biology 241:552–563.

Rossberg, A. 2006b. Some properties of the speciation model for foodweb structure - mechanisms for degree distribution and intervality. Journal of Theoretical Biology 238:401–415.

S., D., and M. Feissel. 2000. Effects of enrichment on threelevel food chains with omnivory. American Naturalist 155:200–218.

Schneider, F., S. Scheu, and U. Brose. 2012. Body mass constraints on feeding rates determine the consequences of predator loss. Ecology Letters 15:436–443.

Smith, D. J., and P. Amarasekare. 2018. Toward a mechanistic understanding of thermal niche partitioning. American Naturalist 191:E57–E75.

Stouffer, D., and J. Bascompte. 2011. Compartmentalization increases food-web persistence. Proceedings of the National Academy of Sciences of the USA 108:3648–3652.

Stouffer, D., J. Camacho, R. Guimerà, C. Ng, and L. L. Amaral. 2005. Quantitative patterns in the structure of model and empirical food webs. Ecology 86:1301–1311.

Suweis, S., F. Simini, J. Banavar, and A. Maritan. 2013. Emergence of structural and dynamical properties of ecological mutualistic networks. Nature 500:449–452.

Thébault, E., and C. Fontaine. 2010. Stability of ecological communities and the architecture of mutualistic and trophic networks. Science 329:853–856.

Tilman, D. 1982. Resource Competition and Community Structure. Princeton University Press.

Wagner, A. 2005. Robustness, evolvability, and neutrality. FEBS Letters 579:1772–1778.

Watts, D., and S. Strogatz. 1998. Collective dynamics of ‘small-world’ networks. Nature 393:440–442.

Williams, R., and N. Martinez. 2000. Simple rules yield complex foodwebs. Nature 404:180–183.

Williams, R. 2008. Success and its limits among structural models of complex food webs. Journal of Animal Ecology 77:512–519.

